# FAZ assembly in bloodstream form *Trypanosoma brucei* requires kinesin KIN-E

**DOI:** 10.1101/2022.12.22.521263

**Authors:** Anna C. Albisetti, Robert L. Douglas, Matthew D. Welch

## Abstract

*Trypanosoma brucei*, the causative agent of African sleeping sickness, uses its flagellum for movement, cell division, and signaling. The flagellum is anchored to the cell body membrane *via* the flagellar attachment zone (FAZ), a complex of proteins, filaments, and microtubules that spans two membranes with elements on both flagellum and cell body sides. How FAZ components are carried into place to form this complex is poorly understood. Here, we show that the trypanosome-specific kinesin KIN-E is required for building the FAZ in bloodstream-form parasites. KIN-E is localized along the flagellum with a concentration at its distal tip. Depletion of KIN-E by RNAi rapidly inhibits flagellum attachment and leads to cell death. A detailed analysis reveals that KIN-E depletion phenotypes include failure in cytokinesis completion, kinetoplast DNA mis-segregation, and transport vesicle accumulation. Together with previously published results in procyclic form parasites, these data suggest KIN-E plays a critical role in FAZ assembly in *T. brucei*.

## Introduction

*Trypanosoma brucei* ssp. are unicellular flagellated parasites endemic to Sub-Saharan Africa that cause human African trypanosomiasis (also known as sleeping sickness) [1] and Nagana disease in cattle [2]. These parasites are transmitted by the bite of infected tsetse flies, proliferate extracellularly in human blood, and eventually reach the central nervous system [3]. *T. brucei* escapes the host adaptive immune system by rapidly replacing its entire surface coat of variable surface glycoproteins (VSG) and by removing bound antibodies [4–6].

To adapt to the different environments encountered during its complex life cycle, *T. brucei* undergoes major cytoskeletal rearrangements as it transitions from procyclic trypomastigote (procyclic form or PCF) in the tsetse fly gut, to epimastigote in the salivary glands, and back to the bloodstream trypomastigote form (bloodstream form or BSF) within the bloodstream of the mammalian host [7]. The parasite relies primarily on its microtubule cytoskeleton, which is composed of sub-pellicular microtubules, the microtubule quartet (MtQ), and the flagellar axoneme consisting of 9 microtubule doublets surrounding a central pair of microtubules. In particular, the flagellum functions in movement, coordination of cytokinesis, and as a sensory organelle [8]. The *T. brucei* axoneme, together with the associated paraflagellar rod (PFR) [9], are contained within the flagellar membrane and connected to the cell body *via* the flagellum attachment zone (FAZ) [10]. The flagellum originates inside the cell body from the basal body (BB), which physically connects it to the mitochondrial DNA (kinetoplast DNA or kDNA) via the tripartite attachment complex (TAC) [11,12]. The flagellum then exits the cell from a membrane invagination called flagellar pocket (FP), which is the sole site depleted of sub-pellicular microtubules, and therefore, the unique site where endocytosis and exocytosis take place [13]. At the flagellum exit point, a cytoskeletal structure called the flagellar pocket collar (FPC) tightens the FP around the flagellar membrane [14,15]. Once outside the FP, the flagellum connects to the cell body via the FAZ, a complex of microtubules, filaments, and proteins that spans and anchors the flagellar and cell body membranes [10]. On the cell body side, the FAZ consists of the FAZ filament and the MtQ. On the extracellular side, transmembrane proteins including FLA1 and FLA1BP connect the two membranes [16,17]. On the flagellar side, FAZ connectors such as FLAM3 link the axoneme to the adhesion region [18]. Because of the functions of the flagellum and FAZ, downregulating components can have dramatic effects on cell physiology. For example, in PCF cells, downregulation of FLAM3 drastically reduces FAZ length and leads to a transition into the epimastigote form [19,20]. In BSF, FLAM3 depletion leads to flagellar detachment and a severe cytokinesis defect, highlighting differences in functions depending on the life cycle stage [20]. FAZ components are numerous, and several play essential roles in regulating cell length, organelle positioning, and cell division [21,22].

The functions of the microtubule cytoskeleton require the activity of kinesin motor proteins, which transport cargoes such as protein complexes, organelles, and chromosomes along microtubules, and regulate microtubule dynamics [23]. The genome of *T. brucei* encodes approximately 50 kinesin-like proteins, many of which are divergent at the level of primary amino acid sequence [24]. Some of these kinesins are classified in families that are unique to kinetoplastids, and some so called orphan-kinesins cannot be classified within any known kinesin subfamily from other species [25,26]. The orphan and kinetoplastid-specific kinesin KIN-E (Tb927.5.2410) [27] was shown in PCF parasites to play an essential role in maintaining trypomastigote morphology and targeting the FAZ component FLAM3 to the flagellum for FAZ assembly [28]. However, the role of KIN-E in the medically-relevant BSF cells has remained undetermined.

In this work, we characterize the localization and function of KIN-E in BSF parasites. We show an essential role for KIN-E in cell survival and FAZ assembly. We also describe morphological phenotypes induced by KIN-E depletion, including impaired cytokinesis and kDNA segregation. Moreover, we find KIN-E is important for FLAM3 localization, suggesting FLAM3 is a cargo of KIN-E in BSF parasites.

## Results

### KIN-E localizes along the flagellum with a concentration at the flagellar tip

To examine the function of KIN-E in BSF parasites, we first sought to localize the KIN-E protein. We raised a polyclonal antibody in rabbits that recognizes the C-terminus of the protein (aa 1105 – 1339) (Fig S1A). By western blotting the antibody detected the recombinant protein expressed in bacteria (Fig S1B-D; tagged with glutathione-S-transferase (GST)) and recognized a single band of approximately 150 kDa in *T. brucei* cell extract, corresponding to the annotated molecular mass of 149,644 Da (Fig S1E). By immunofluorescence microscopy in BSF cells, the anti-KIN-E antibody stained the lengths of old and new flagellum with a signal enrichment at the tip of the new flagellum (Fig 1). Occasion signal enrichment could also be seen on the tip of the old flagellum, and non-specific cytoplasmic staining was also sometimes observed. KIN-E signal was seen at the tips of both short (Fig 1B) and long (Fig 1C) new flagella, suggesting it selectively tracks the growing flagellum tip.

**Fig 1.**
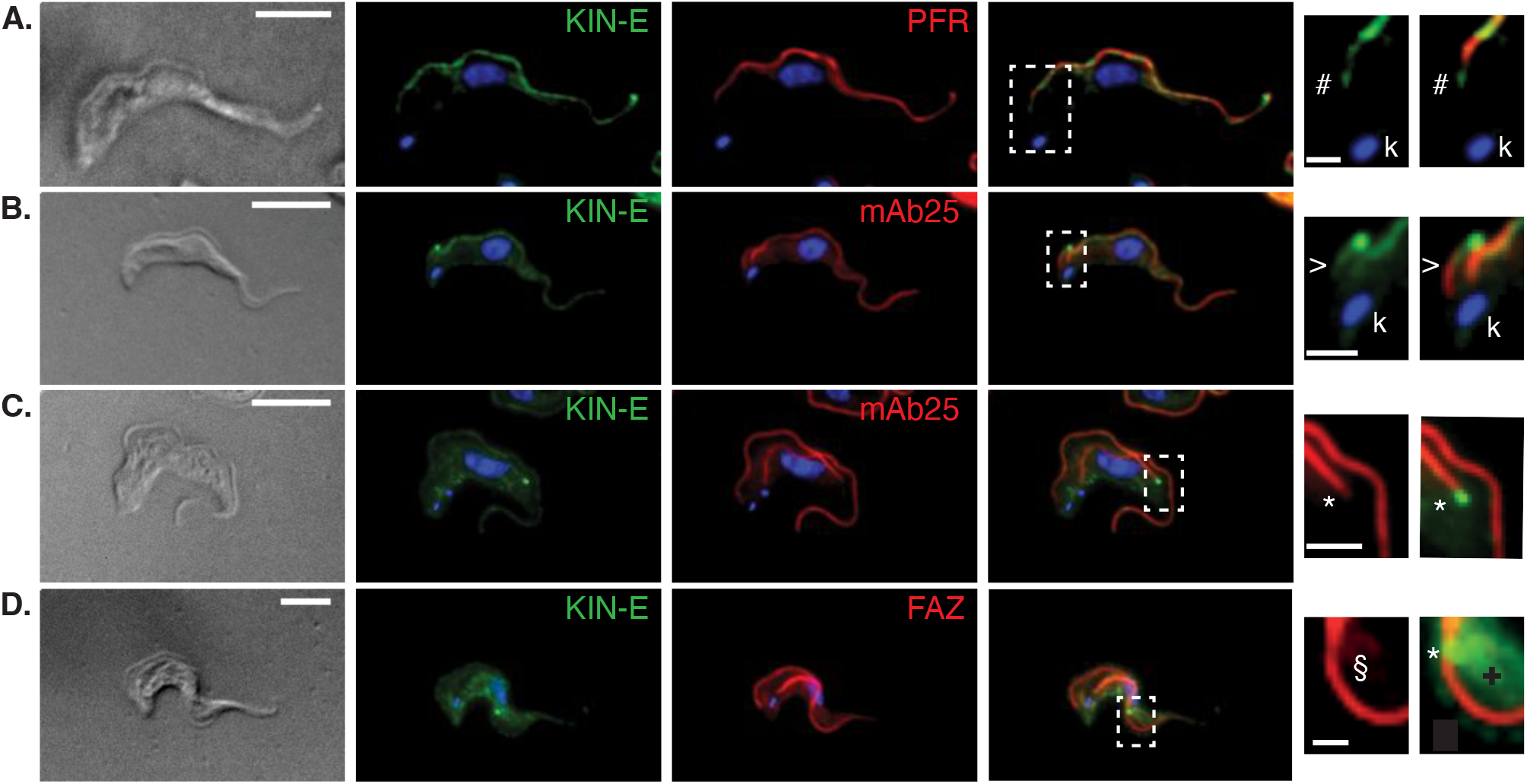
KIN-E localizes along the flagellum with a concentration at the flagellar tip. **(A-D)** *T. brucei* BSF 90-13 cells visualized by differential interference contrast (DIC) microscopy, or immunofluorescence microscopy using anti-KIN-E (green) and also stained for DNA (blue, DAPI) and the following markers (red): **(A)** L8C4, a PFR marker; **(B-C)** mAb25, an axonemal marker; and **(D)** L3B2, a FAZ marker. White dashed boxes show magnified views of: **(A,B)** the proximal end of the flagellum, with (A) # indicating a comparison of the proximal end of the KIN-E signal with the L3B2; and (B) > a comparison of the proximal end of the KIN-E signal with mAb25; **(C)** KIN-E staining at the new flagellum tip (*) compared to mAb25 at the anterior (distal) end of the new flagellum; **(D)** Brightness enhanced image of KIN-E on an old flagellum adjacent to the FAZ signal on the cell body side (§), with cytoplasmic staining also evident (+). KIN-E staining (*) can also be seen at the anterior flagellar tip in (A) and on the anterior tip of very short (B) or already longer new flagella (C & D). (k) indicates the kDNA. Scale bars 5 *μ*m (left) and 1.25 *μ*m (right).

To gain more detailed insight into KIN-E localization at the proximal end of the flagellum and along its length, we compared its distribution with that of known flagellar markers. In cells stained for KIN-E and a marker of the PFR (L8C4) [29], KIN-E extended more towards the proximal end of the flagellum (nearer to the BB) than the PFR (Fig 1A). On the other hand, in cells stained for KIN-E and the axonemal marker mAb25 [30], the mAb25 signal extended more towards the proximal end of the flagellum than the KIN-E signal (Fig 1B). We also stained for KIN-E and a marker of the cell body side of the FAZ (L3B2) [29], and found that KIN-E and the FAZ followed parallel paths, but did not overlap (Fig 1D). Together, these results indicate that, in addition to the flagellar tip localization, KIN-E staining initiates at the proximal end of flagella distal to the start of the axoneme but proximal to the start of the PFR, likely corresponding to a region near the FPC.

### KIN-E is essential for bloodstream form *T. brucei* survival

To study the function of KIN-E in BSF parasites, we generated a stable and tetracycline-inducible RNAi cell line (KIN-E^RNAi^). We monitored cell growth of the parental cell line (90-13), and of uninduced and induced KIN-E^RNAi^ cells. Upon KIN-E RNAi induction, we observed growth arrest starting at 24 h post induction (hpi), followed by cell death (as measured by a decrease in cell number) within 72 hpi (Fig 2A). The efficiency of KIN-E knock-down was confirmed by western blotting. At 24 hpi, KIN-E expression dropped below 20% of that in uninduced cells, and at 48 hpi KIN-E was not detected (Fig. 2B). KIN-E signal was also monitored by immunofluorescence microscopy at 16 hpi (Fig 2Ci) and 24 hpi (Fig 2Cii) in cells counterstained for the flagellar axoneme (mAb25). KIN-E signal was diminished along the flagellum after 16 h and 24 h (although some residual signal remained at flagellar tip; compare with Fig 1). These data indicate that KIN-E expression is efficiently downregulated by RNAi, resulting in premature cell death. Thus, KIN-E is essential for BSF parasite survival.

**Fig 2.**
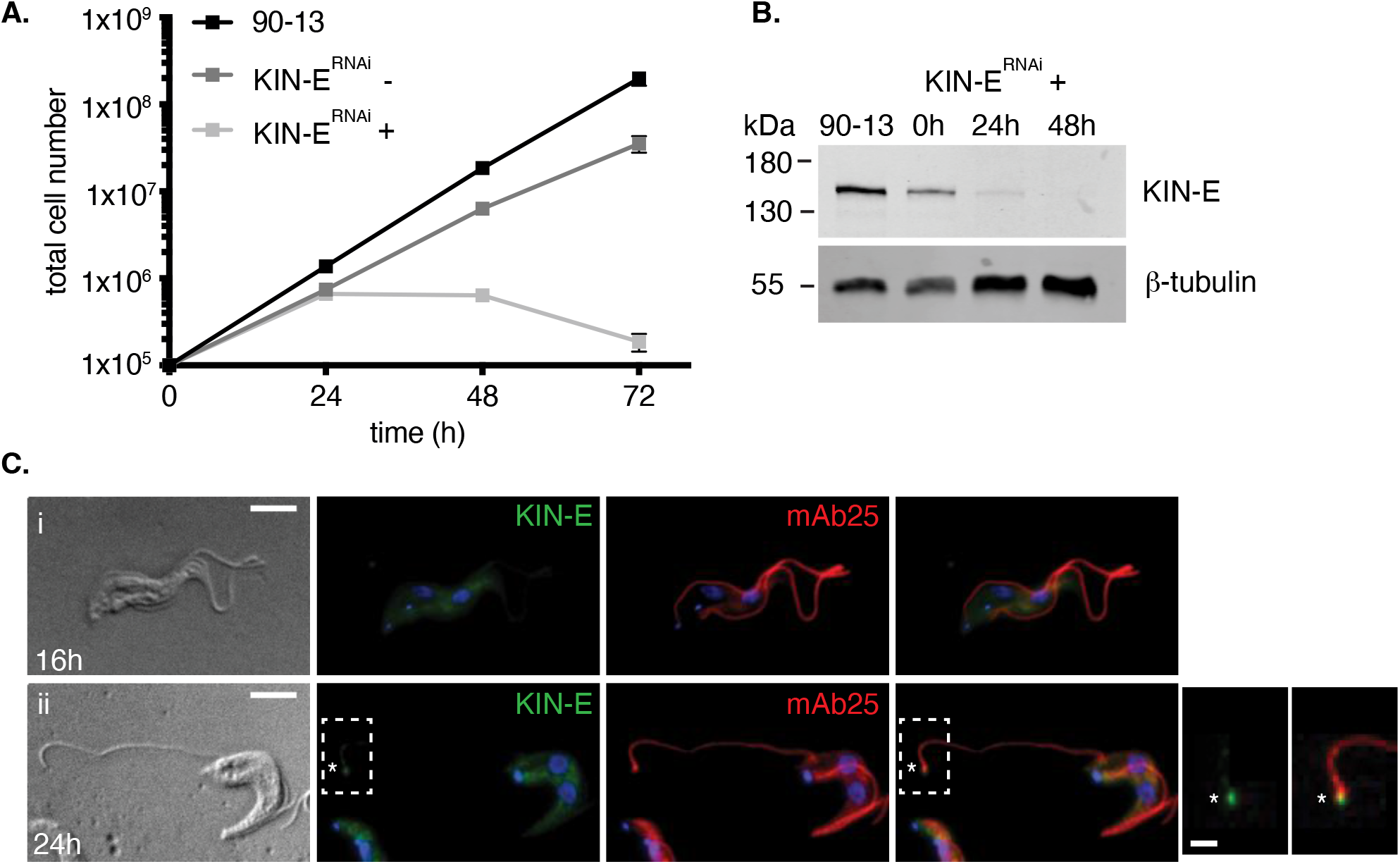
KIN-E is essential for bloodstream form *T. brucei* survival. **(A)** Growth curve of the parental cell line (90-13) compared to uninduced (−) and induced (+) KIN-E^RNAi^ cells. Error bars show SE, n=3 independent experiments. **(B)** KIN-E protein expression during RNAi induction analyzed by western blotting with anti-KIN-E antibody and anti-β-tubulin antibody as a loading control. Equal cell numbers (5×10^6^ cells) were loaded per lane. **(C)** KIN-E^RNAi^ cells visualized by DIC or immunofluorescence microscopy at (i) 16 hpi or (ii) 24 hpi using anti-KIN-E (green) and mAb25 (red) as an axonemal marker. Residual new flagellar tip signal in (ii) is marked with an asterisk (*). Scale bars 5 *μ*m (left) and 1.25 *μ*m (right).

### KIN-E is required for the attachment of the newly synthetized flagellum, but not for flagellum biogenesis or beating

In observing the morphology of KIN-E^RNAi^ cells during the first 24 hpi, we noted that many had partially detached new flagella, although cells otherwise looked relatively normal (Fig 2C, 3A). We quantified the percentage of cells with a detached flagellum and found that it increased from ~20% at 16 hpi to ~50% at 24 hpi (Fig 3A). We hypothesized that failure in flagellum attachment was caused by improper FAZ assembly. To assess FAZ integrity, we stained KIN-E^RNAi^ cells for a FAZ marker (L3B2) at 16 hpi and 24 hpi. We observed that FAZ staining was interrupted at the location where the new flagellum detached from the cell body, although the FAZ staining associated with the old flagellum was unaffected (Fig 3Bi, 3Bii). This suggests that new flagella become detached, whereas old flagella remain attached. Detached new flagella and attached old flagella were also visible by transmission electron microscopy (TEM) at 16 hpi (Fig 3C). These findings suggest that KIN-E plays a role in FAZ assembly, and that downregulating KIN-E function causes defects in this process that result in detachment of the new flagellum.

**Fig 3.**
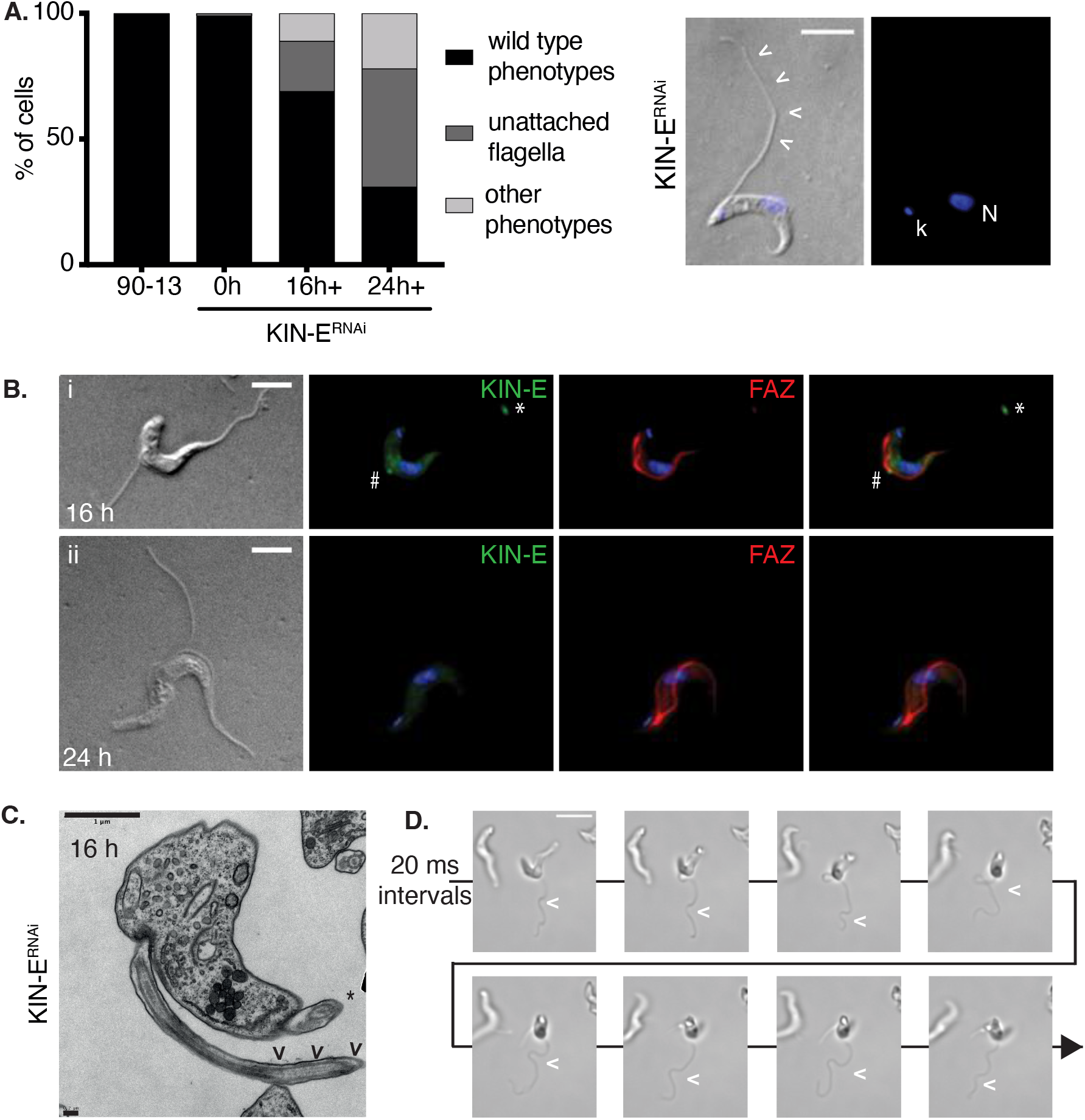
KIN-E is required for the attachment of the newly synthetized flagellum. **(A)** Graph of the percentage of KIN-E^RNAi^-uninduced or -induced cells with detached flagella or other abnormal phenotypes at 16 hpi and 24 hpi compared with parental cell line 90-13 >200 cells per condition, n=3 independent experiments. Image on right shows an example of a detached flagellum (arrowheads) in KIN-E^RNAi^ at 16 hpi visualized with DIC. DNA was stained with DAPI (blue). Scale bar 5 μm. **(B)** KIN-E^RNAi^ cells visualized by DIC or immunofluorescence microscopy at (i) 16 hpi or (ii) 24 hpi, using anti-KIN-E (green) and L3B2 (red) as a FAZ marker. Residual KIN-E signal is visible at the point of detachment of a new flagellum (#) and at the tip of old flagellum (*). **(C)** TEM image of KIN-E^RNAi^ cells at 16 hpi. Arrowheads (<) indicate the detached new flagellum. The attached old flagellum on the same cell is marked with an asterisk (*). Upper left scale bar 1 μm; lower left scale bar 0.2 μm. **(D)** Video frames of KIN-E^RNAi^ cells at 24 hpi, acquired every 20 ms. Flagellar wave in a detached flagellum is indicated with an arrowhead. Scale bar 10 *μ*m. Also see Supplementary Video 3.

We also investigated whether motility of the detached new flagellum was affected by KIN-E depletion. We recorded movies of 90-13 parental cells as well as KIN-E^RNAi^ cells with detached flagella at 24 hpi (Fig 3D and Supplementary Videos 1 - 4). We observed active beating of detached as well as attached flagella (Video 4 features an unusual abnormal dividing cell with two attached and two detached beating flagella). Altogether, this suggests that KIN-E is essential for flagellar attachment, but not for flagellar biogenesis or beating.

#### KIN-E is critical for proper cytokinesis

Previous studies reported that defects in FAZ assembly often lead to a failure in cytokinesis [31,32]. To determine the effect of KIN-E depletion on cell cycle progression and cytokinesis in the BSF, we performed a comprehensive phenotypic analysis, focusing on quantifying nuclear DNA (N) and kinetoplast DNA (kDNA or K) content (Fig 4A). We divided the cell population in three categories: cells with wild-type DNA content (1K1N, 2K1N, 2K2N) and attached flagella; cells with wild-type DNA content and detached flagella; and cells with abnormal DNA content, with or without detached flagella. The latter category included: multinucleated cells, indicating failed cytokinesis (>2K2N) (Fig 4A, Bi); cells with one kDNA but two nuclei, indicating failed kDNA replication/division (1K2N) (Fig 4Bii); and other phenotypes (1K0N, 0K1N, and indeterminate DNA content (?K?N)). As expected, almost all 90-13 parental cells had wild-type DNA content and attached flagella (Fig 4A). In contrast, KIN-E^RNAi^ cells at 16 hpi and 24 hpi showed an increasing percentage of cells with abnormal DNA content, including 20% multinucleated cells at 24 hpi (Fig 4A). Multinucleated cells were also observed by TEM (Fig 4C). Thus, KIN-E is critical for efficient cytokinesis, likely as a consequence of disturbed FAZ assembly.

**Fig 4.**
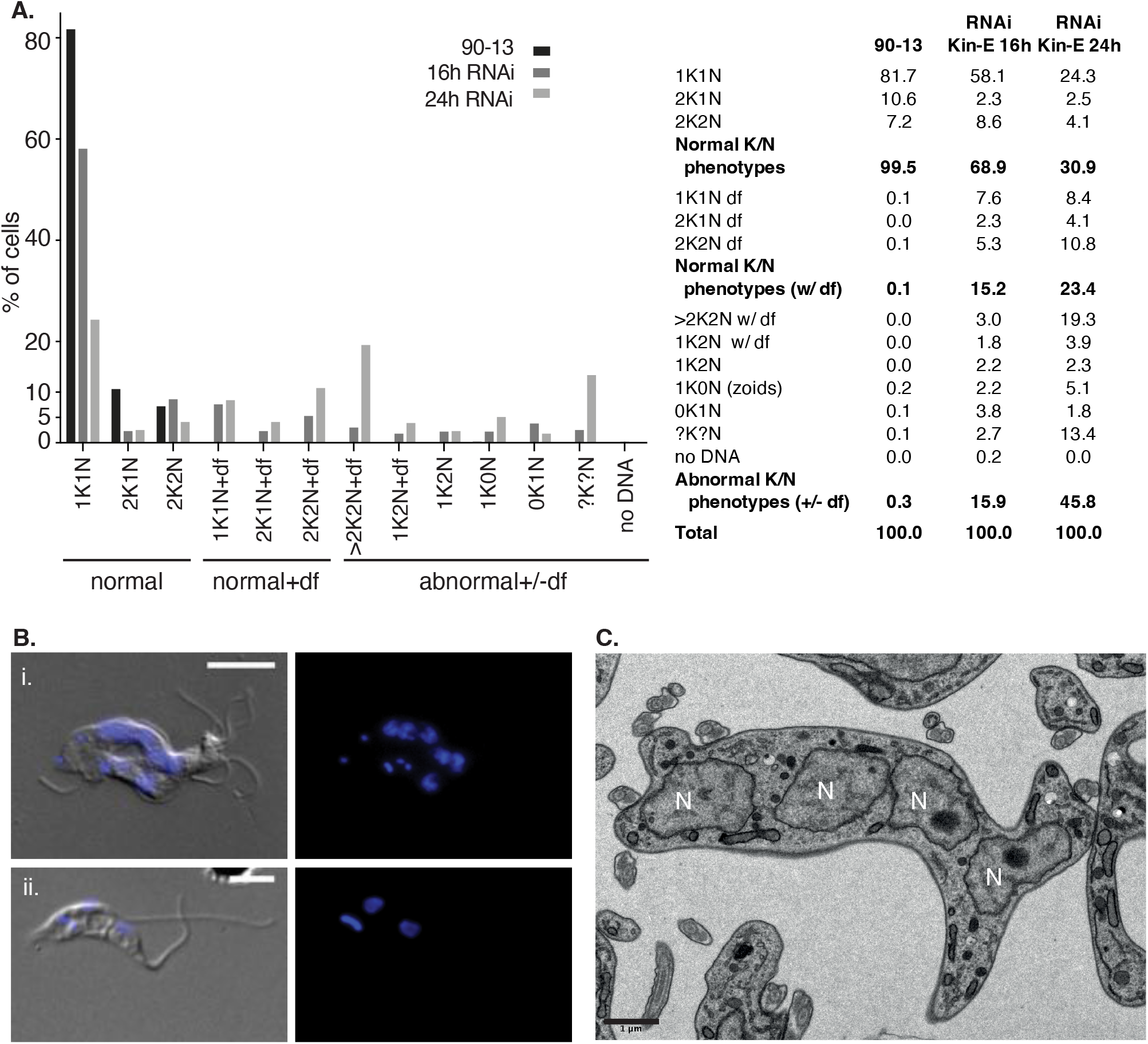
KIN-E depletion causes a failure in cytokinesis. **(A)** Graphical and tabular representations of phenotypic counts with a focus on DNA content in KIN-E^RNAi^ cells at 16 hpi and 24 hpi compared with the parental cell line 90-13. The categories were defined as follows: normal kinetoplast (K) and nucleus (N) phenotypes (1K1N, 2K1N, 2K2N); normal K/N phenotypes with detached flagella (+df); abnormal K/N phenotypes with or without detached flagella (+/-df), including cells with a single kDNA but two nuclei (1K2N), anucleated cells (1K0N, zoids), multinucleated cells (>2K2N), and other phenotypes (?K?N). n>200 cells per cell line. **(B)** Examples of (i) a multinucleated cell (>2K2N) and (ii) a 1K2N cell. Nuclei and kinetoplasts were stained with DAPI (blue); cell bodies and flagella were visualized with DIC. Scale bars 5 μm. **(C)** Thin section TEM micrograph of KIN-E^RNAi^ population induced for 24 hpi, showing a cell with multiple nuclei (N). Scale bar 1 μm.

#### KIN-E influences kDNA segregation

As mentioned above, we observed that 1K2N cells represented up to 7% of KIN-E^RNAi^ cells at 16 hpi and 24 hpi in (Fig 4A, Fig 5Ai). We sought to further test whether these were defective in kDNA duplication or kDNA segregation. Because the kDNA is physically attached to the flagellum [11], we first imaged flagella (by staining the axoneme with mAb25) and FPC structures (by staining for BILBO1; [33]) in 1K2N cells (Fig 5Aii). We found that each kDNA was attached to two separated FPCs and two flagella, suggesting that the kDNA had duplicated but not segregated.

**Fig 5.**
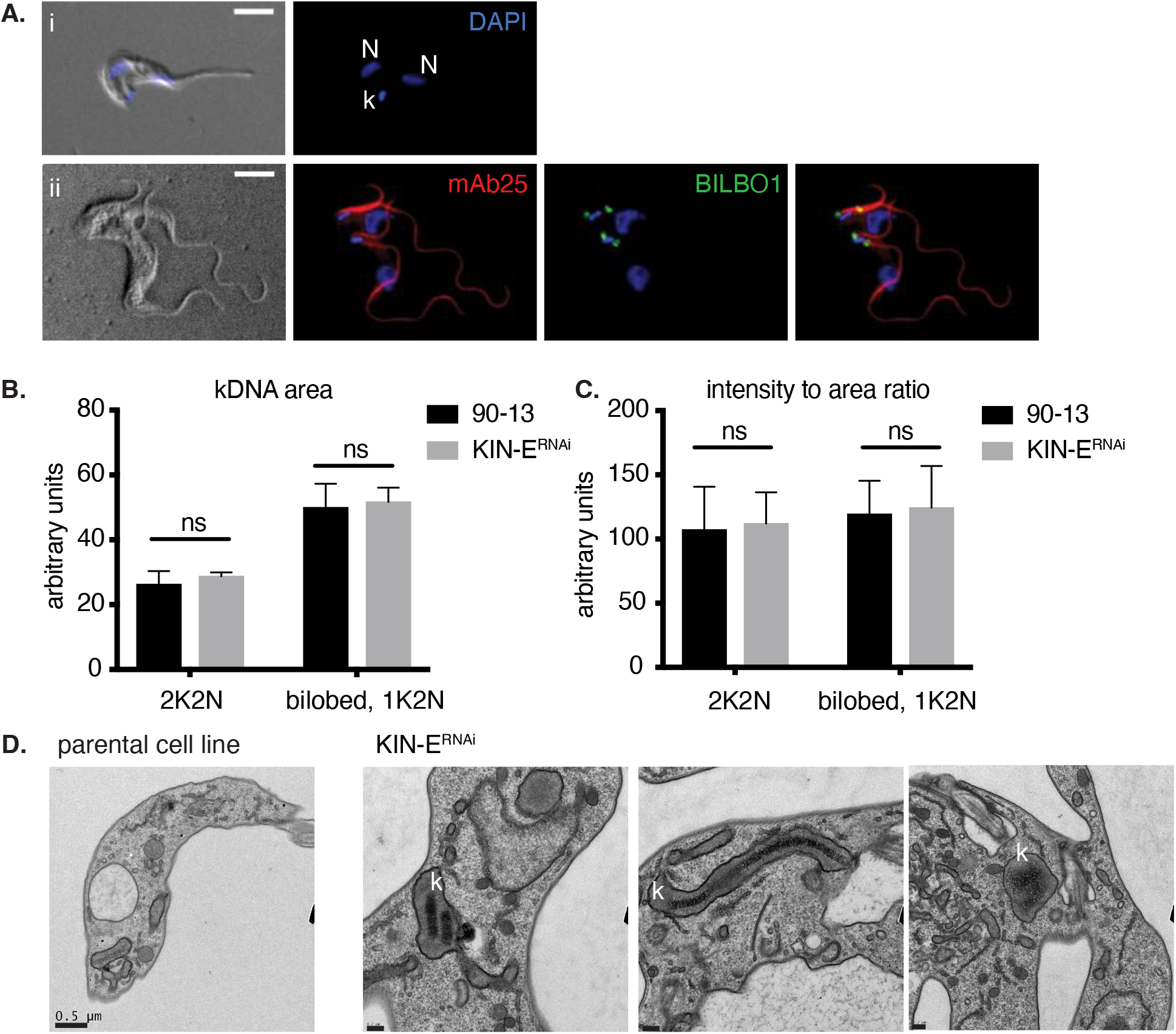
KIN-E influences kDNA segregation. **(A)** KIN-E^RNAi^ cells at 24 hpi, showing (i) a 1K2N cell with a single enlarged kDNA and two nuclei, and (ii) a 2K2N cell with 4 flagella stained with mAb25 antibody (red), and 4 FPCs stained with anti-BILBO1 antibody (green). Scale bars 5 μm. **(B)** kDNA area and **(C)** ratio of kDNA signal intensity / area comparing single kDNAs in KIN-E^RNAi^ induced 2K2N cells versus parental 90-13 2K2N cells, as well as the single kDNA (1K2N) in KIN-E^RNAi^-induced cells versus the bilobed-shaped kDNA in parental 90-13 cells. Quantification of 100 cells per condition, n=3 independent experiments. Statistical comparisons between strains were performed using a t-test, ns = non-significant. **(D)** Thin section TEM micrographs of parental cells and KIN-E^RNAi^ cells at 16 hpi, 16 hpi, and 24 hpi, showing different abnormal kDNA (k) configurations. Scale bars 0.5 μm for the parental cell line, and 0.2 μm for KIN-E^RNAi^.

To further test for kDNA duplication/segregation, we separately measured the area of each single DAPI-stained kDNA in parental 90-13 and induced KIN-E^RNAi^ 2K2N cells, as well as each kDNA in induced KIN-E^RNAi^ 1K2N cells. The mean area of kDNA in KIN-E^RNAi^ 1K2N cells was twice as large as each single kDNA in 2K2N cells (Fig 5B). Additionally, the area of kDNA in 1K2N KIN-E^RNAi^ was comparable to the dividing bilobed-kDNA [34] in the parental cell line (Fig 5B). We also measured kDNA signal intensities and calculated the intensity:area ratio. On average, the kDNAs in 1K2N KIN-E^RNAi^ and dividing bilobed-kDNA parental cells were both twice as intense and twice as large as the single kDNAs in 2K2N cells, resulting in a comparable intensity:area ratio (Fig 5C). Finally, by TEM we observed in 7 of 72 sections that the kDNA was in an atypical configuration, appearing as two compacted or one elongated disc, or as a disorganized structure (Fig 5D). From these data, we conclude KIN-E does not influence mitochondrial DNA duplication, but does influence mitochondrial DNA segregation.

#### KIN-E is important for vesicular trafficking near the flagellar pocket

In the vicinity of the FP in TEM sections we also observed an increased number of vesicles in the KIN-E^RNAi^-induced cells compared with parental cells (Fig 6A). At 16 hpi KIN-E^RNAi^ cells had an average of 6 ± 0.4 vesicles, and at 24 hpi KIN-E^RNAi^ cells had an average of 7 ± 0.5 vesicles, whereas parental 90-13 cells had 3 ± 0.5 vesicles per FP (Fig 6B). The diameter of vesicles was indistinguishable in parental versus induced KIN-E^RNAi^ cells, and averaged 115 ± 2 nm (Fig 6C, D). In BSF *T. brucei*, endocytosis occurs at the FP via large clathrin-coated vesicles (135 nm in diameter) containing variable surface glycoproteins (VSG) [35,36]. We observed clathrin-coated vesicles in the process of being internalized (Fig 6C). The shape and size of the observed vesicles in the parental cell line, uninduced KIN-E^RNAi^ cells, and induced KIN-E^RNAi^ population were similar to endocytic vesicles after shedding their clathrin-coat. The luminal side of the vesicles showed an electron-dense material of the same thickness as the coat on the cell surface and flagellar membrane (Fig 6C), suggesting this material is VSG. For these reasons, we speculate that these vesicles near the FP are endocytic vesicles.

**Fig 6.**
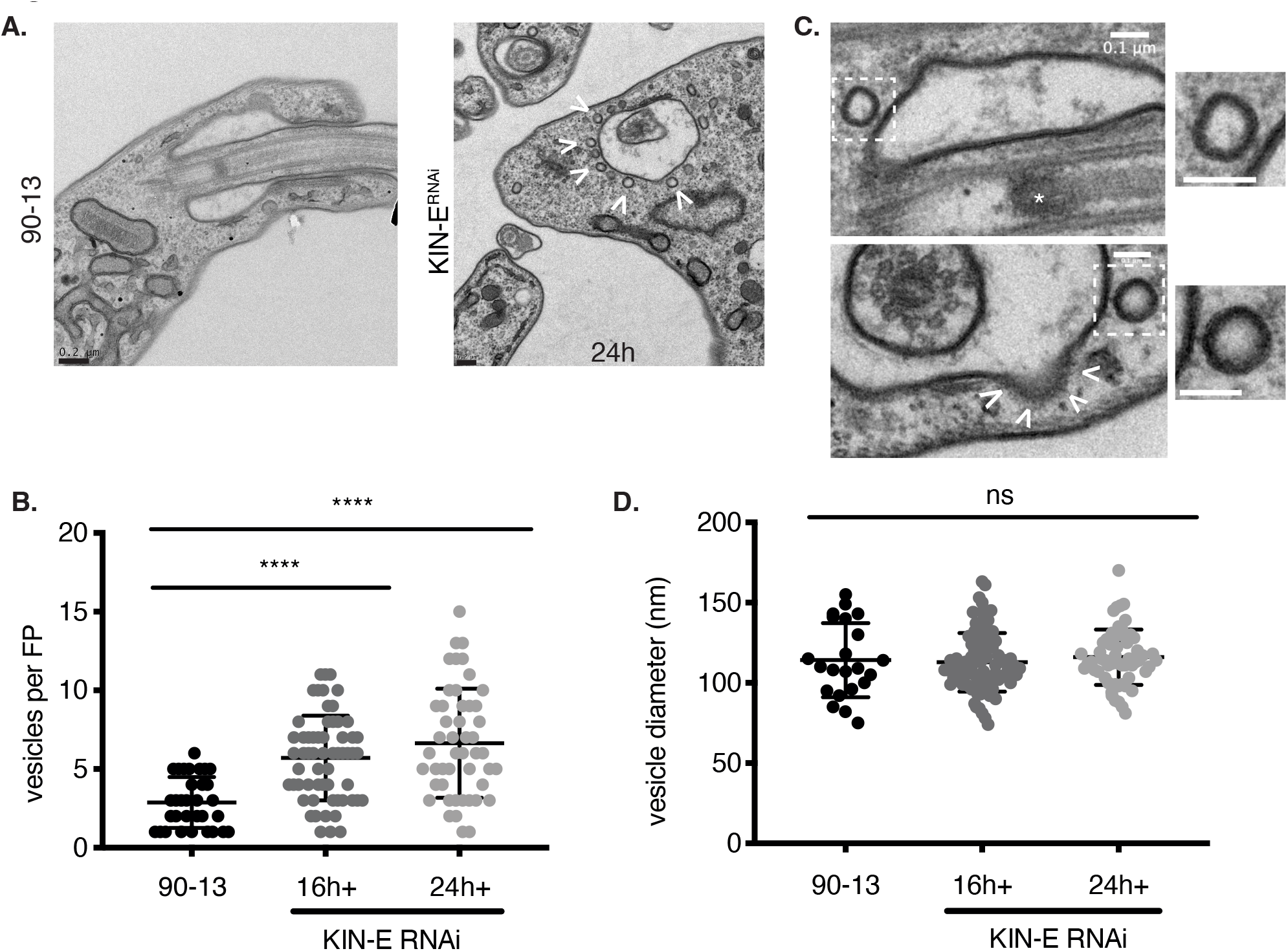
KIN-E downregulation leads to vesicle accumulation near the FP. **(A)** Thin section TEM micrographs comparing the FP region of the parental cell line 90-13 with that of a KIN-E^RNAi^ at 24 hpi. Scale bar 0.2 μm. **(B)** Number of vesicles per FP for the parental cell line 90-13 and KIN-E^RNAi^ cells at 16 hpi and 24 hpi. n>200 vesicles per condition. Bars show group mean as well as top and bottom quartiles. Pairwise statistical comparisons were performed using a t-test, **** p<0.0001. **(C)** Thin section TEM micrographs show vesicles near the FP in KIN-E^RNAi^ cells. White dashed boxes show magnified views of individual vesicles. The upper panel shows a longitudinal cross section of the flagellum showing the basal plate (*). The lower panel shows a transverse cut around the basal plate of the flagellum. Arrowheads mark a clathrin-coated vesicle budding from the FP. Scale bar 0.1 μm. **(D)** Vesicle diameter was measured from TEM micrographs using ImageJ. n=165 total vesicles measures. Pairwise statistical comparisons were performed using a t-test, ns = non-significant.

### *KIN-E* is required for FLAM3 localization to the new flagellar tip

Our observation that KIN-E depletion disrupts FAZ formation suggests that it functions to transport cargoes that are known components of the FAZ [19,20,31,37–40]. One potential KIN-E cargo is FLAM3, which was previously identified as a FAZ component that is located on the flagellar side, accumulates at the new flagellar tip, and depending on the cell cycle stage, shows weak or no localization to the old flagellar tip [19,20]. Because FLAM3 has a comparable localization to KIN-E at the new flagellar tip in PCF cells [20], we tested whether FLAM3 exhibited a similar localization in BSF cells. We endogenously tagged FLAM3 at its C-terminus with a 10x myc tag (FLAM3_myc_) within the KIN-E^RNAi^ cell line background. The growth of uninduced cells was unaffected by the expression of FLAM3_myc_ (Fig 7A). We localized FLAM3_myc_ within the old and the new flagellum by immunofluorescence microscopy and observed signal enrichment at the new flagellar tip (Fig 7B), but no signal at the old flagellar tip (Fig 7B). We next tested whether KIN-E depletion had an effect on FLAM3 localization by examining the distribution of FLAM3 in induced KIN-E^RNAi^ cells at 24 and 48 hpi (Fig 7B). As described above for KIN-E^RNAi^ cells (Fig 2A), induced KIN-E^RNAi^ FLAM3_myc_ cells died within 72 hpi and showed detached new flagella. Interestingly, although FLAM3 was at the tip of new flagella at 0 hpi, at 24 hpi, and 48 hpi, FLAM3 was confined to the proximal end of the new flagellum (Fig 7B), between the kDNA and the origin of the PFR. The old flagellum still showed weak FLAM3 staining in flagella (labeled with mAb25) at 24 hpi, whereas flagellar FLAM3 signal was nearly absent at 48 hpi. As control, we used KIN-E antibody to confirm that KIN-E was properly depleted at 48 hpi (Fig 7B), while FLAM3 was present at the proximal end of the flagellum and diffuse within the cytosol (Fig 7B). The altered FLAM3 localization suggests that, in BSF parasites, KIN-E is required for FLAM3 transport into the flagellum.

**Fig.7.**
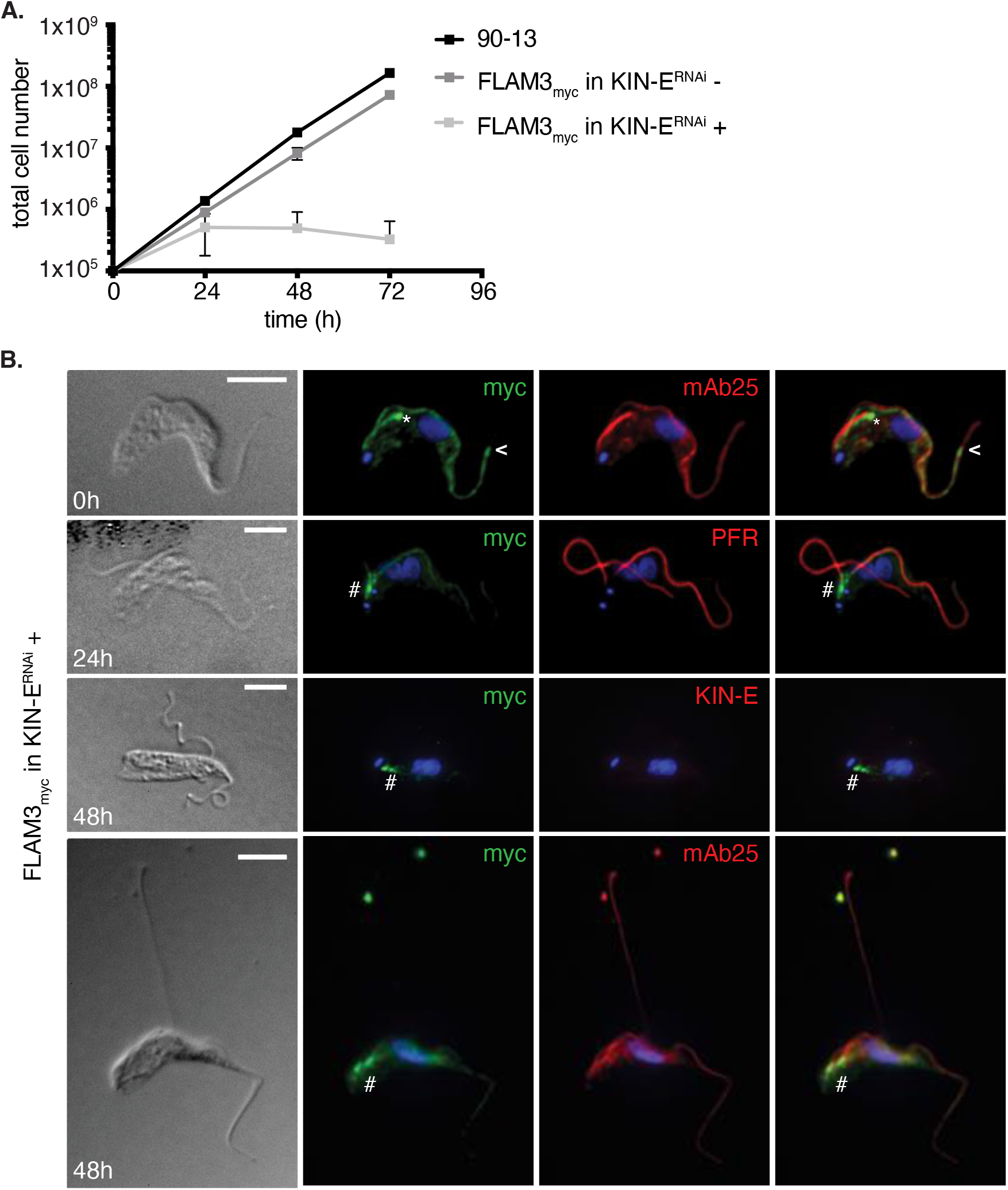
FLAM3 is a KIN-E cargo. **(A)** Growth curve of the parental cell line 90-13 compared with FLAM3_myc_ in KIN-E^RNAi^-uninduced (−) and -induced (+) cells. **(B)** FLAM3_myc_ KIN-E^RNAi^ cells visualized by DIC or immunofluorescence microscopy. Immunofluorescence analysis of endogenously expressed FLAM3-10myc in KIN-E^RNAi^ uninduced (0 hpi) and induced cells at 24 hpi and 48 hpi using anti-myc to detect FLAM3 (green), and mAb25 or PFR (red) as a flagellar marker. Asterisk (*) marks the bright FLAM3 signal at the new flagellar tip, arrowheads (<) show the point where FLAM3 signal stops near the distal end of the flagellum, hash tag (#) indicates FLAM3 signal accumulation at the proximal end of the flagellum. The two bright dots near the top of l the bottom row 48 hpi image are likely flagellar debris from dead cells. Scale bar 5 μm.

## Discussion

We investigated the localization and function of the kinetoplastid-specific kinesin KIN-E in BSF *T. brucei*. We found that KIN-E is localized within the flagellum, with an enrichment at the distal tip of growing new flagella. We further observed that KIN-E is essential for cell survival of *T. brucei* BSF parasites. KIN-E is necessary for attachment of the newly synthesized flagellum and biogenesis of the FAZ. In the absence of KIN-E expression, cytokinesis fails, as does kDNA segregation in a subset of cells. Our work establishes important roles for KIN-E in BSF trypanosomes.

Our phenotypic analysis in BSF confirms and extends the characterization of KIN-E function in PCF trypanosomes [28]. In PCF cells, KIN-E is also localized to the flagellum and enriched at the flagellar tip [28,41]. However, in BSF cells, we observe significant KIN-E signal enhancement at the tips of new flagella, whereas in PCF cells it is seen at the tips of both old and new flagella [28,41]. Furthermore, in PCF cells, KIN-E is important for normal growth rate [28], although we find it is essential for viability in BSF cells. KIN-E depletion in PCF cells also induces the repositioning of the kDNA and the production of epimastigote-like cells [28], outcomes we do not observe in BSF cells.

A key function of KIN-E in both PCF and BSF cells is in flagellar attachment. KIN-E depletion in PCF cells causes a failure in attachment of newly synthetized flagella [28], similar to what we observe in BSF cells. In both PCF and BSF parasites, cells with detached new flagella contain a full length old FAZ filament but a short new FAZ filament, suggesting premature termination of FAZ synthesis. The old FAZ filament is not affected by KIN-E depletion, suggesting that once this component of the FAZ on the cell body side is synthesized, KIN-E is no longer required for its maintenance. However, on the flagellar side, FAZ maintenance could be affected by KIN-E depletion. Furthermore, KIN-E is not required for flagellar length, which *in T. brucei* is controlled by the intraflagellar transport (IFT) machinery [31,42], as it is in other organisms [43]. In further support of the notion that KIN-E is not required for flagellar function, we found that detached flagella continue to beat in KIN-E-depleted BSF cells.

One cargo of KIN-E in PCF cells is FLAM3 [28], a component of the FAZ on the flagellar side [19]. Upon KIN-E depletion in PCF cells, FLAM3 becomes diffusely localized within the cytosol [28]. Our work suggests that FLAM3 is also a cargo of KIN-E in BSF cells. Interestingly, upon KIN-E depletion in BSF, we find that FLAM3 accumulates at the proximal end of the flagellum in an area that may overlap with the transition zone, which has been described as the “gate” that controls transport of components into the flagellum [44,45]. There may also be other KIN-E cargoes, for example, the proteins ClpGM6 and FAZ27, which colocalize and interact with FLAM3 on the flagellar side of the FAZ [20,37]. Future research will establish their interactions and the mechanism of FAZ assembly.

We observed a second major phenotype upon KIN-E depletion in BSF cells, which is the accumulation of multinucleated and multi-flagellated cells, as well as a small population of zoids, indicative of a failure in cytokinesis. Normal flagellum structure and function is important for cytokinesis in *T. brucei*, and in BSF *T. brucei*, flagellar defects cause a failure in cytokinesis and cell inviability [46–50]. RNAi knockdown of IFT proteins also produces defects in flagellum construction and causes impaired cytokinesis [31,51,52]. *T. brucei* cytokinesis also requires flagellar attachment, and loss of FAZ proteins like FAZ1 [39], FLA1 [38], FAZ10 [40], or FLAM3 [19,20] leads to defects in cytokinesis. Thus, cytokinesis failure and cell inviability in KIN-E-depleted BSF cells is likely a consequence of defects in FAZ synthesis and flagellar attachment.

We found that depletion of KIN-E also causes other phenotypes. For example, a subset of KIN-E-depleted cells (<10%) fail in segregating their kDNA. In *T. brucei*, kDNA is physically linked to the flagellar BBs *via* the TAC complex [11,12], and its segregation is orchestrated by the movements of the BBs [53]. However, the forces involved in this process remain mostly unknown. It has been suggested that the MtQ could drive BB movement and consequently kDNA segregation [53]. Moreover, a properly formed FAZ is indispensable for BB segregation and cell division [31]. Failure in kDNA segregation may therefore be a consequence of impaired FAZ formation. A separate phenotype observed upon KIN-E depletion is the accumulation of vesicles around the FP that contain VSG in their lumen and thus are likely endocytic vesicles that have shed their clathrin coat. Why these vesicles accumulate remains mysterious. Further investigations could elucidate how KIN-E impacts kinetoplast segregation and vesicular trafficking in the vicinity of BBs and FP.

The further study of KIN-E and its cargos could help elucidate how the FAZ is assembled and maintained in *T. brucei*. Moreover, KIN-E is a kinetoplastid-specific kinesin with orthologs in the related organisms *Trypanosoma cruzi* and *Leishmania spp*. [27]. The fact that KIN-E is essential for parasite survival makes it a potential drug target. Kinesin inhibitors have been identified with promising drug-like properties and have been tested as anti-cancer drugs [54–56]. This raises the possibility that KIN-E-targeting drugs could be developed to treat human and animal African trypanosomiasis, as well as Chagas disease and Leishmaniasis.

## Materials and Methods

### Cell lines, growth conditions and transfections

*T. brucei* strain 427 90-13 BSF cells [57] were cultured at 37°C in HMI-9 medium [58] supplemented with 10% fetal bovine serum (FBS, Atlanta Biologicals), 2.5 μg/ml G418 (InvivoGen), and 5 μg/ml hygromycin (InvivoGen).

Plasmid transfections into 90-13 cells were performed using the Amaxa Nucleofector™ system (Lonza) with program X-001 as described previously [59], and with Tb-BSF buffer (90 mM Na_2_HPO_4_, 5 mM KCl, 0.15 mM CaCl_2_, 50 mM HEPES, pH 7.3) [60]. Stable cell lines were selected by culturing cells in medium containing 2.5 μg/ml phleomycin (InvivoGen) and/or 10 μg/ml blasticidin (InvivoGen). Expression of double-stranded RNA was induced by adding 1 μg/ml tetracycline (Sigma-Aldrich) to the culture medium.

### Plasmid construction

For RNAi silencing of KIN-E expression in *T. brucei*, we PCR-amplified a DNA segment (bp 1798 - 2333) of the KIN-E gene (Tb927.5.2410) from genomic DNA isolated from *T. brucei* 90-13 cells using the Qiagen DNeasy Blood and Tissue kit. Suitability of this segment for RNAi was confirmed using the online tool RNAit [61]. The purified PCR product was inserted by standard ligation into XhoI and HindIII sites within the pZJM vector [62] containing the phleomycin resistance (*ble*) gene, between two opposing T7 promoters. A total of 10 μg of the plasmid was linearized with NotI for transfection into bloodstream form *T. brucei* 90-13 cells (carried out as described above).

To generate a *T. brucei* strain expressing endogenous C-terminally myc-tagged FLAM3 (FLAM3_myc_), we used the long primer PCR transfection method described previously [63]. We used the pPOTv7 plasmid DNA as template for PCR amplification, which contained coding sequences for the 10x myc tag and blasticidin resistance cassette. Transfection was carried out as described above.

For protein expression and purification of glutathione-S-transferase (GST) fused to the C-terminal domain of KIN-E in *E. coli* (amino acids 1105 – 1339; GST-KIN-E^C-ter^), bp 3313 – 4017 of the corresponding gene were amplified by PCR from genomic DNA, and cloned into BamHI and NotI restriction sites within the pGEX-4T-1 plasmid (GE Healthcare), such that the gene was in frame with an N-terminal GST tag (pGEX-4T-1-GST-KIN-E^C-ter^). All plasmid sequences were confirmed by DNA sequencing at the UC Berkeley DNA Sequencing Facility.

### Protein expression and purification for antibody production

The plasmid pGEX-4T-1-GST-KIN-E^C-ter^ was transformed into *E. coli* strain BL21. Bacteria grown in 1 l lysogeny broth (LB) with 100 *μ*g/ml ampicillin were induced with 1 mM isopropyl-β-D-thio-galactoside (IPTG) for 2.5 - 4 h at 37°C. Bacteria were centrifuged (4000 x g for 20 min at 4°C) and resuspended in cold lysis buffer (50 mM Tris, pH 8, 50 mM NaCl, 5 mM EDTA, 0.2% Triton-X 100, 1 mM β-mercaptoethanol, 150 μm PMSF, and 1 μg/ml final volume each of leupeptin, pepstatin, and chymostatin (LPC) protease inhibitor mix). Cells were lysed on ice using a Branson Digital Sonifier 450 (at level 6 for 1 min (10 s on, 10 s off, x 3) + 30 min rest, x 3-4 cycles). Cell lysate supernatant was loaded on a Glutathione Sepharose 4B column (GE Healthcare). The column was washed with TBS (20 mM Tris, pH 7.5, 150 mM NaCl) with 5 mM EDTA, 0.1% Triton-X 100, 1 mM β-mercaptoethanol, 150 μm PMSF, and LPC. Washed bound protein was then eluted with elution buffer, 50 mM Tris pH 8.0, plus 10 mM reduced glutathione, into 10 fractions. The column fraction with the greatest protein concentration were then subjected to gel filtration chromatography on a Superdex 75 column equilibrated with 50mM Tris, pH 8.0, plus 150 mM NaCl. Fractions containing GST-KIN-E were collected and stored at −80°C.

### Antibody production and purification

Two rabbits were immunized (Covance Inc.) with 1.5-2 mg of purified GST-KIN-E^C-ter^ protein, according to Covance’s 118 d protocol. KIN-E^C-ter^-specific antibodies were affinity purified as follows. Purified GST-KIN-E^C-ter^ fusion protein from pooled gel filtration fractions was cross-linked to Affi-gel 15 beads (BioRad) in MOPS, pH 7.0, with 1M KCl, and the beads were quenched with ethanolamine HCl, pH 8.0, at 4°C. Serum from immunized rabbits was loaded onto the column, and the column was then washed with 20 mM Tris, pH 7.6, 0.5 M NaCl and 0.2% Triton X-100. Purified bound antibody was eluted with 200 mM glycine pH 2.5, 150 mM NaCl, and 10 × 300 *μ*l fractions were collected into fraction tubes containing 50 *μ*l 1 M Tris-HCl, pH 8.0. To test for antibody specificity by western blotting, bacteria were cultivated at 37°C, induced for 2 h 30 min with 1 mM IPTG, harvested, boiled in sample buffer, and subjected to SDS-PAGE.

### Western blotting

Western blotting was performed using standard methods as described previously [64], with the exception that proteins were transferred to nitrocellulose membranes (Genesee Scientific, Prometheus #84-875). Primary antibodies were used at the following dilutions: rabbit anti-KIN-E 1:2,000; rabbit anti-GST 1:1,000 (Welch lab); mouse anti-β-tubulin E7 1:10,000 (Developmental Studies Hybridoma Bank, University of Iowa). Secondary antibodies used were goat anti-rabbit AF790 (ThermoFisher A11367) and goat anti-mouse AF680 (ThermoFisher A21058), both diluted at 1:10,000. Images were taken using the Odyssey imaging system (Li-Cor Biosciences).

### Fluorescence microscopy

For immunofluorescence microscopy, parental and TbKIN-E^RNAi^-uninduced and -induced cells in exponential growth phase were harvested for 5 min at 1,000 x g at room temperature and washed once in Voorheis’ modified PBS (vPBS; 8 mg/ml NaCl, 0.22 mg/ml KCl, 2.27 mg/ml Na_2_HPO_4_, 0.41 mg/ml KH_2_PO_4_, 15.7 mg/ml sucrose, 1.8 mg/ml glucose). The cells were resuspended in 1% paraformaldehyde (PFA) in vPBS and incubated for 2 min on ice. Cells were centrifuged for 5 min at 2,000 x g, resuspended in vPBS, and settled on slides for 10 min. Slides were incubated for 30 min in −20°C methanol and rehydrated for 10 min in PBS (8 mg/ml NaCl, 0.2 mg/ml KCl, 1.44 mg/ml Na_2_HPO_4_, 0.24 mg/ml KH_2_PO_4_). Cells were incubated for 1 h with primary antibodies diluted in PBS (rabbit anti-KIN-E 1:1,000; mouse mAb25 1:10 [30]; mouse L8C4 (PFR) 1:10 [29]; mouse L3B2 (FAZ) 1:10 [29]; and rabbit anti-BILBO1 1:4,000 [33]). Cells were washed 3 times in PBS and incubated in a dark moist chamber for 1 h with secondary antibodies, all at a 1:200 dilution in PBS (anti-rabbit AF488 (ThermoFisher A11008), anti-rabbit AF564 (ThermoFisher A11036), anti-mouse AF488 (ThermoFisher A11001), anti-mouse AF568 (ThermoFisher A11004), anti-rat AF488 (ThermoFisher A21208)). Kinetoplasts and nuclei were stained with DAPI (10 μg/ml) for 4 min. Slides were mounted with ProLong Gold antifade reagent (ThermoFisher P36930). Images were acquired with a Zeiss AxioImager microscope, equipped with a Hamamatsu Orca 03 camera, using iVision software version 4.5.6r4 (BioVision Technologies), and analyzed with Fiji ImageJ version 1.51 [65].

For DAPI staining of nuclei and kinetoplasts, parental and TbKIN-E^RNAi^-uninduced and -induced cells (at 0 h, 16 h, and 24 h after induction with 1 μg/ml tetracycline) in mid-log phase were harvested and washed once in vPBS. The cells were fixed in 1% paraformaldehyde in vPBS for 2 min on ice. Cells were centrifuged 5 min at 2,000 x g, resuspended in vPBS, and settled onto glass slides. Cells were permeabilized 30 min in −20°C methanol, rehydrated with PBS, and stained with DAPI (1 μg/ml) for 3 min. Slides were mounted with ProLong Gold antifade. Images were acquired using a Zeiss AxioImager microscope and Hamamatsu Orca 03 camera with the same exposure time between samples, using iVision software version 4.5.6r4, and analyzed and quantified with Fiji ImageJ version 1.51.

### Live cell imaging

90-13 cells and TbKIN-E^RNAi^ were deposited onto glass bottom dishes 24 h following induction with 1 μg/ml tetracycline. Images were acquired at 20 frames/s with an Olympus IX71 microscope, equipped with an optiMOS™ sCMOS camera, using Micro-manager software [66] and analyzed with Fiji ImageJ version 1.51.

### Electron microscopy

For TEM, 90-13 and TbKIN-E^RNAi^ cells induced for 16 h or 24 h with 1 μg/ml tetracycline were collected in mid-log phase and fixed for 30 min in HMI-9 with 2% glutaraldehyde at room temperature, and incubated overnight at 4°C. Cells were pelleted for 10 min at 1,000 x g, resuspended in 2% very low gelling-point agarose (Sigma A5030) in water, pelleted again, and incubated for 15 min on ice. Agarose was cut into small pieces, rinsed twice in 0.1 M sodium cacodylate buffer (NaO_2_As(CH_3_)_2_), and incubated for 1 h in 1% osmium tetroxide (OsO_4_) in 0.1 M sodium cacodylate buffer with 1.6% potassium ferricyanide (pH 7.2). After three washes with 0.1 M sodium cacodylate buffer (pH 7.2), cells were dehydrated in successively higher concentrations of acetone (35%, 50%, 70%, 80%, 95%, 100%, and 100%) for 10 min incubations each. Cells were then incubated in acetone:resin (Eponate 12P) 2:1 for 30 min, 1:1 for 30 min and 1:2 for 1 h, followed by incubation in pure resin for 72 h. Cells were centrifuged for 10 min in a benchtop microfuge and moved into a new tube containing pure resin with accelerant benzyl dimethylamine (BDMA) for 30 min, and then for 4 h. Resin was left to polymerize overnight at 60°C in silicon molds or flat bottom capsules. Ultra-thin sections were cut with an Ultracut E microtome (Reichert Jung) to approximately 70 nm thick. Sections were loaded on formvar-coated mesh or slot grids (Electron Microscopy Sciences, G100-Cu, S2010-NOTCH) and stained with 2% uranyl acetate and lead citrate. Samples were visualized on Tecnai 12 transmission electron microscope with a UltraScan®1000XP CCD Camera and with the Gatan Digital micrograph software, and processed with Fiji ImageJ version 1.51.

### Bioinformatic analysis

Distinct prediction software was used to identify domains within the KIN-E protein. The motor domain (aa 14-336) was identified using Pfam [67]. The ARM domains (aa 480-653) were identified using SMART (Simple Modular Architecture Research Tool) [68]. The CalpainIII-like domains (aa 713-997) were identified as in [28] by alignment of the m-calpain domain III-like domains (mCL#1 and mCL#2) of KIN-E with the domain III of the human m-calpain protein (PBD code: 1KFU). The coiled-coil domain (aa 1171-1213) was identified using SMART.

### Statistical analysis

The statistical parameters and significance are reported in the figure legends. Statistical analyses were performed using GraphPad PRISM v.8.

## Acknowledgments

We acknowledge an important contribution from Haiming Wu, Ph.D., who purified the KIN-E protein as well as anti-KIN-E antibodies. We wished to include Haiming Wu as an author, but have been unable to contact him for several years. Due to the bioRxiv policy that all authors must consent to submission, we regret we are unable to include him in the author list. We also thank the following individuals: Mélanie Bonhivers and Derrick Robinson for anti-BILBO1 and mAb25 antibodies; Keith Gull for L3B2 (FAZ) and L8C4 (PFR) antibodies; Samuel Dean for the pPOTv7 plasmid; Denise Schichnes and Steven Ruzin of the UC Berkeley RCNR Biological Imaging Facility for assistance with light microscopy; and Reena Zalpuri and Danielle Jorgens of the UC Berkeley Electron Microscope Lab for TEM training. Finally, we thank Neil Fischer for proofreading the manuscript. Funding was provided by RLD, including through a family trust.

## Author contributions

AA, RLD, and MDW designed the research; AA and RLD performed research; AA analyzed data; AA wrote the original draft; AA, RLD, and MDW reviewed and edited the manuscript; RLD and MDW acquired funding.

## Supplementary Figure and Video Legends

**Fig S1.**
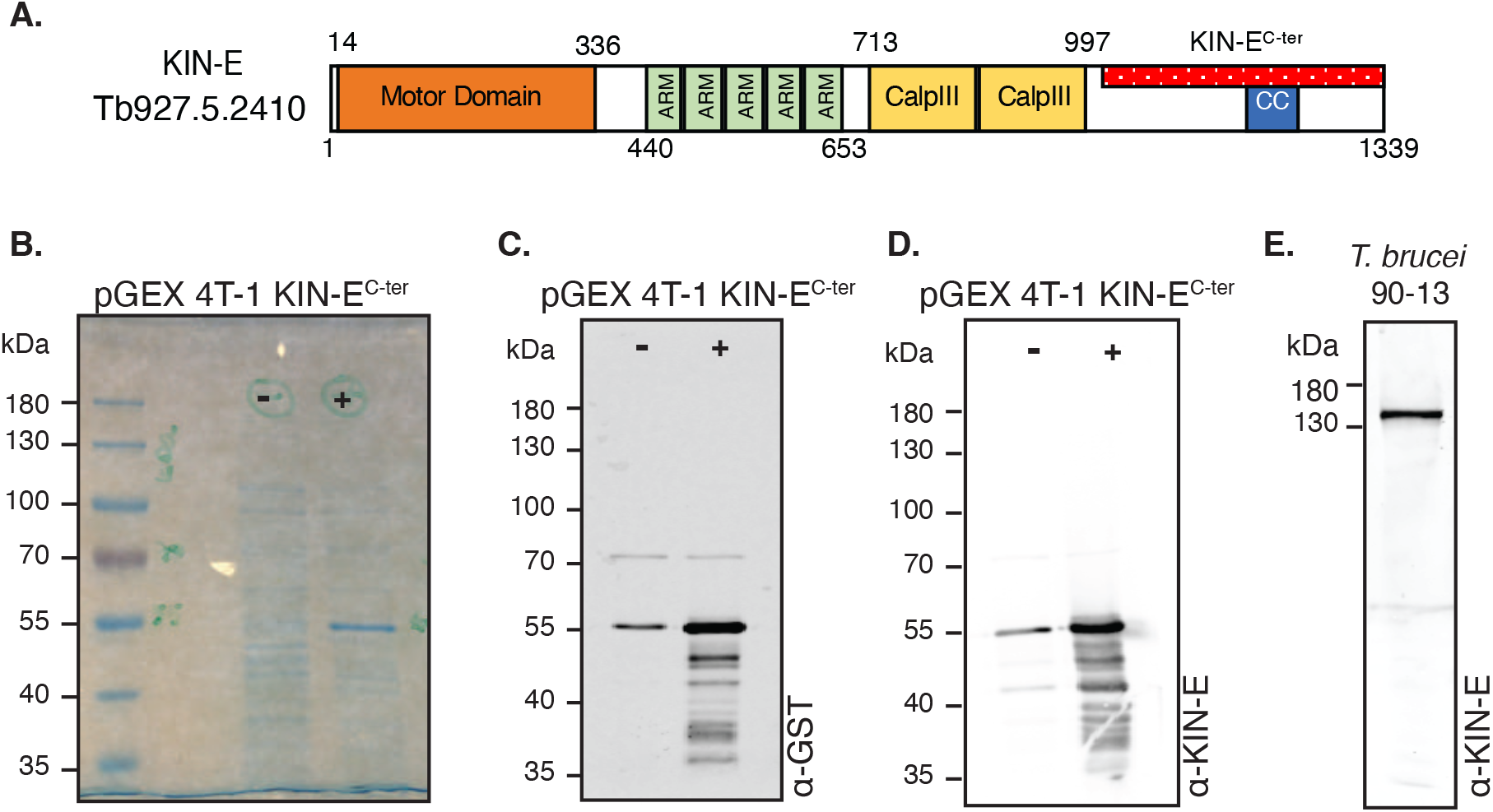
Specificity of the rabbit anti-KIN-E antibody tested by western blotting. **(A)** Schematic representation of KIN-E (Tb927.5.2410), comprising the motor domain (orange), the armadillo repeats (ARM, green), the two CalpainIII-like domains (CalpIII, yellow), the short coiled-coil domain (CC, blue) and the KIN-E^C-ter^ (1105-1339 aa, red with dots, used to raise an anti-KIN-E antibody). **(B)** SDS-PAGE gel, stained with SimpyBlue™ SafeStain, showing the expression of GST-KIN-E^C-ter^ in 1 mM IPTG-induced (+) bacteria versus uninduced (−) bacteria. **(C,D)** Western blots of bacterial extracts (from 5×10^8^ cells in each lane) either uninduced or induced to express GST-KIN-E^C-ter^ and probed with (C) anti-GST or (D) anti-KIN-E antibodies. **(E)** Western blot of *T. brucei* 90-13 whole cell lysate (from 5×10^6^ *T. brucei* cells) probed with anti-KIN-E antibody, showing a single band at around 150 kDa (KIN-E molecular mass 149 kDa).

**Video 1** Flagellum beating in a 90-13 cell. Frames acquired every 20 ms, plays at 25 frames/sec.

**Video 2** Flagella beating in a dividing 90-13 cell. Frames acquired every 20 ms, plays at 25 frames/sec.

**Video 3** Detached flagellum beating in a KIN-E^RNAi^ cell at 24 hpi. Frames acquired every 20 ms, plays at 25 frames/sec.

**Video 4** Detached flagella beating in a dividing KIN-E^RNAi^ cell at 24 hpi. Frames acquired every 20 ms, plays at 25 frames/sec.

